# Lipid nanoparticle-mediated RNAi targeting of KRAS mutants sensitises colorectal cancer cells to chemotherapy

**DOI:** 10.64898/2026.10.15.759489

**Authors:** Yuhang Wang, Shuquan Su, Kainat Zahra, Rui Sang, Mark P. Molloy, Gyorgy Hutvagner, Wei Deng

## Abstract

KRAS mutations, including G12D, G12V and G13D, are major oncogenic drivers in colorectal cancer (CRC) and are associated with therapeutic resistance. However, approved KRAS inhibitors remain focused on G12C subtype, leaving these prevalent variants without approved targeted therapies. Here, we developed a lipid nanoparticle (LNP)-based RNA interference (RNAi) strategy to target these three KRAS variants using a single small interfering RNA (siRNA) sequence. The developed siRNA-LNPs reduced KRAS mRNA and protein expression in HCT116 (G13D), SW620 (G12V) and LS180 (G12D) cells, but not in KRAS wild-type HT29 cells, suppressing the cellular proliferation, clonogenic survival and migration. Transcriptomic profiling revealed changes in KRAS associated signalling, including suppression of cell-cycle and DNA repair programmes, providing a rationale for combining siRNA silencing with the fluoropyrimidine chemotherapeutic agent 5-fluorouracil (5-FU). The combination produced synergistic effects across all three KRAS-mutant CRC cell models and enhanced the apoptosis in both monolayer cultures and 3D spheroids. Collectively, these findings provide proof of concept for this combination strategy and support its further investigation as a potential therapeutic approach for KRAS-mutant CRC.

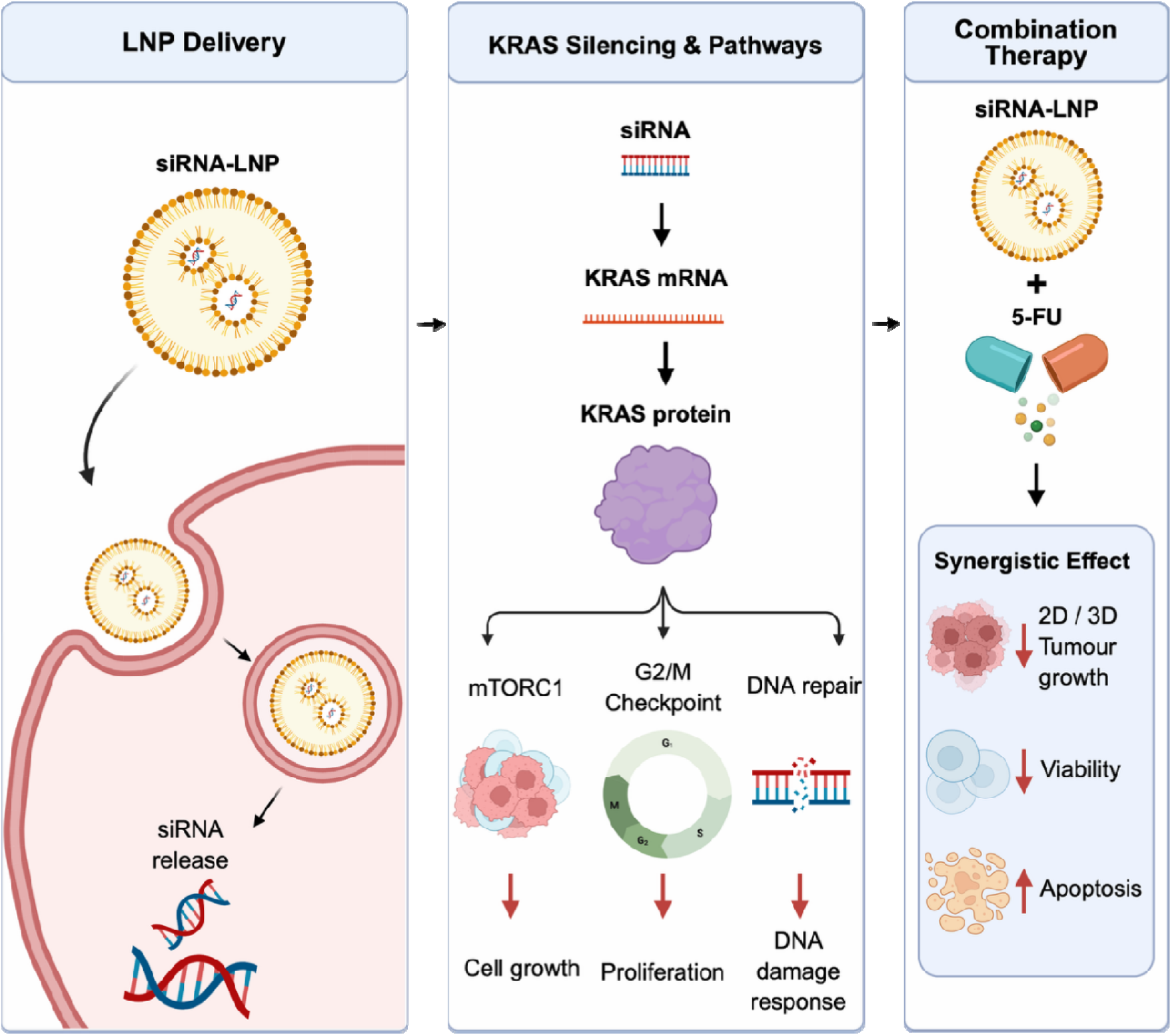

## Introduction

Colorectal cancer (CRC) is the third most commonly diagnosed cancer worldwide and the second leading cause of cancer-related death, with an estimated 2.04 million new cases and 918,000 deaths reported globally in 2024 [1]. Adjuvant fluoropyrimidine-based chemotherapy is the current standard treatment for stage III colon cancer to reduce the risk of recurrence; however, clinical outcomes remain poor in KRAS-mutant CRC[2]. KRAS mutations occur in approximately 40-45% of CRC and are associated with significantly poorer disease-free survival compared with KRAS wild-type disease[3–5]. This poor prognosis is further compounded by limited therapeutic options, as KRAS mutations have been associated with reduced responsiveness to chemotherapy and resistance to anti-EGFR therapies [6, 7].

KRAS has historically been difficult to drug due to its high affinity for GTP/GDP and the lack of drug binding pockets [8]. Recent success in KRAS inhibitor development have predominantly focused on the G12C subtype which can be targeted through the chemically reactive Cys residue, although this subtype accounts for only a small proportion of KRAS mutations in CRC and has shown limited durability due to acquired drug resistance [9–12]. In contrast, the more prevalent KRAS variants in CRC, including G12V, G12D and G13D, remain therapeutically intractable [6]. Therapeutic approaches have therefore also focused on downstream KRAS signalling pathways, including RAF-MEK-ERK and PI3K-AKT-mTOR [13, 14]. However, these strategies are limited by dose dependent toxicities due to other pathway inhibition in normal tissues and signalling feedback loops that allow tumours to escape drug antagonism [13, 14].

RNA interference (RNAi) provides an alternative approach to KRAS suppression through small interfering RNA (siRNA)-mediated, sequence-specific degradation of complementary mRNA transcripts, resulting in reduced expression of the target protein [15, 16]. Because this strategy operates at the transcript level rather than depending on a druggable protein pocket, siRNA-based silencing offers a versatile approach for targeting KRAS gene [16]. Previous studies have used transferrin-targeted cyclodextrin-polymer nanoparticles and DOPC nanoliposomes to deliver KRAS or RAS pathway-targeting siRNAs in preclinical CRC models [17, 18]. However, these approaches were not mutant-selective, potentially limiting their ability to selectively target KRAS-mutant cancer cells and increasing the risk of off-target effects [17]. More recent studies have investigated siRNAs targeting pan-KRAS or G12V mutants, but these approaches have been evaluated predominantly in pancreatic cancer models, with very limited investigation in KRAS-mutant CRC [18, 19]. Collectively, these studies support the potential of RNAi for KRAS suppression, but highlight a gap in the evidence for targeting prevalent KRAS variants in CRC.

In this study, we developed an siRNA-based strategy to selectively target KRAS mutants in CRC cell lines. LNP formulated siRNA suppressed KRAS expression and reduced cellular proliferation, clonogenic survival and migration across KRAS-mutant CRC cell lines. Transcriptomic profiling revealed coordinated changes in KRAS-associated signalling, cell cycle and DNA repair pathways following KRAS silencing. This siRNA-based approach also enhanced the response to 5-fluorouracil in KRAS-mutant cell models, with combinatorial efficacy further demonstrated in 3D tumour spheroids. Together, these findings support the potential of combining KRAS-targeted siRNA silencing with chemotherapy as a therapeutic strategy for KRAS-mutant CRC.

## Result

### Development and characterisation of siRNA-loaded lipid nanoparticles

To develop LNPs suitable for siRNA delivery, key microfluidic fabrication parameters, including lipid composition, buffer, flow rate ratio (FRR), total flow rate (TFR) and purification method, were systematically optimised (Table S1). In our experiment condition, Tris buffer and centrifugal ultrafiltration produced smaller and more uniform nanoparticles than the other conditions tested, while an FRR of 1:1 and TFR of 12 mL/min yielded the smallest particle size and lowest PDI. Based on these results, a DOTAP/DOPE molar ratio of 1:1 was selected for subsequent LNP-siRNA formulations.

Using the optimised formulation conditions, four LNP formulations were prepared: empty LNPs, coumarin-6-loaded LNPs (Cou6-LNP), scrambled siRNA-loaded LNPs (siScr-LNP), and KRAS-targeting siRNA-loaded LNPs (siKRAS-LNP) (Figure 1A). DLS analysis showed that all formulations had particle sizes below 150 nm, with empty LNPs measuring approximately 76 nm and cargo-loaded LNPs ranging from 110-120 nm (Figure 1B). Zeta potential decreased from +55 mV for empty LNPs to +36 mV following siRNA encapsulation, consistent with electrostatic interactions between the cationic lipids and negatively charged siRNA (Figure 1C). Encapsulation efficiencies exceeded 90% for all cargo-loaded formulations, with siKRAS-LNPs achieving approximately 93% (Figure 1D). Cryo-TEM further confirmed a relatively uniform spherical morphology, with particle sizes consistent with the DLS measurements (Figure 1G). The colloidal stability of siKRAS-LNPs was subsequently evaluated under different pH and cell culture conditions. Particle size and PDI remained largely stable over 24 h in PBS at pH 5-7 and in serum-free McCoy’s 5A medium. In medium containing 10% FBS, PDI increased modestly without a substantial change in particle size (Figure 1E, F). These results demonstrate that siKRAS-LNPs maintained their physicochemical characteristics under the tested conditions.

**Figure 1.**
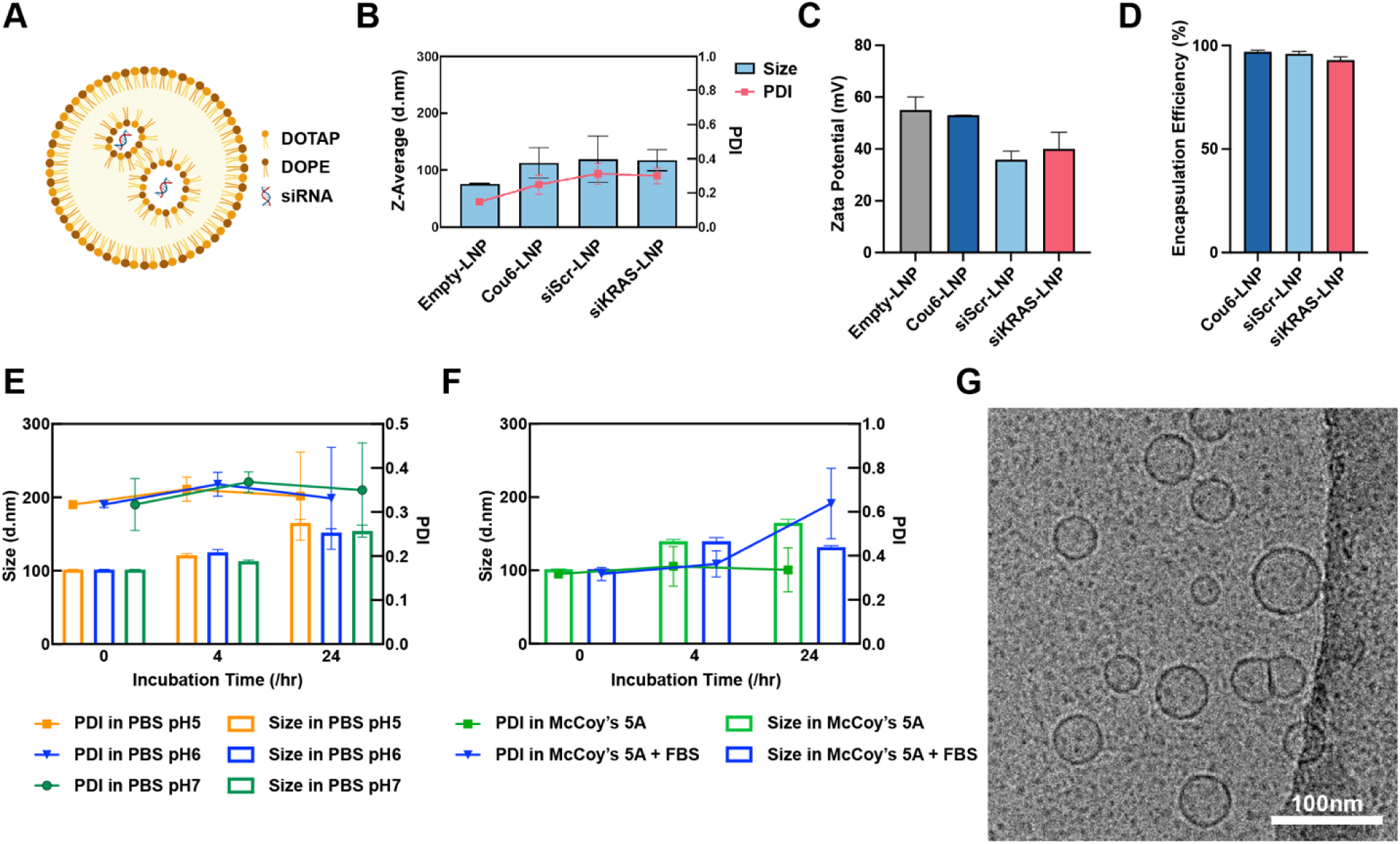
Physicochemical characterisation of LNPs. (A) Schematic of siRNA-LNP composition. (B) Hydrodynamic diameter and polydispersity index (PDI) of empty LNP, Cou6-LNP, siScr-LNP, and siKRAS-LNP. (C) Zeta potential of LNP formulations. (D) Encapsulation efficiency of cargo-loaded LNPs. (E) Stability of siKRAS-LNPs in PBS at pH 5-7 over 24 h. (F) Stability of siKRAS-LNPs in McCoy’s 5A medium with or without 10% FBS over 24 h. (G) Representative cryo-TEM image of empty LNPs. Scale bar, 200 nm. Data in (B–F) are presented as mean ± SD from n = 3 independent experiments.

### Cellular uptake of LNPs

Cellular uptake was evaluated using Cou6-LNPs to assess the internalisation of LNPs by KRAS-mutant colorectal cancer cells. HCT116, SW620 and LS180 cells were incubated with Cou6-LNPs for 2 or 4 h and analysed by confocal microscopy. Intracellular coumarin-6 fluorescence was detected in all three cell lines, with fluorescence distributed throughout the cytoplasm surrounding the DAPI-stained nuclei (Figure 2A), indicating cellular uptake of the Cou6-LNP-associated cargo.

**Figure 2.**
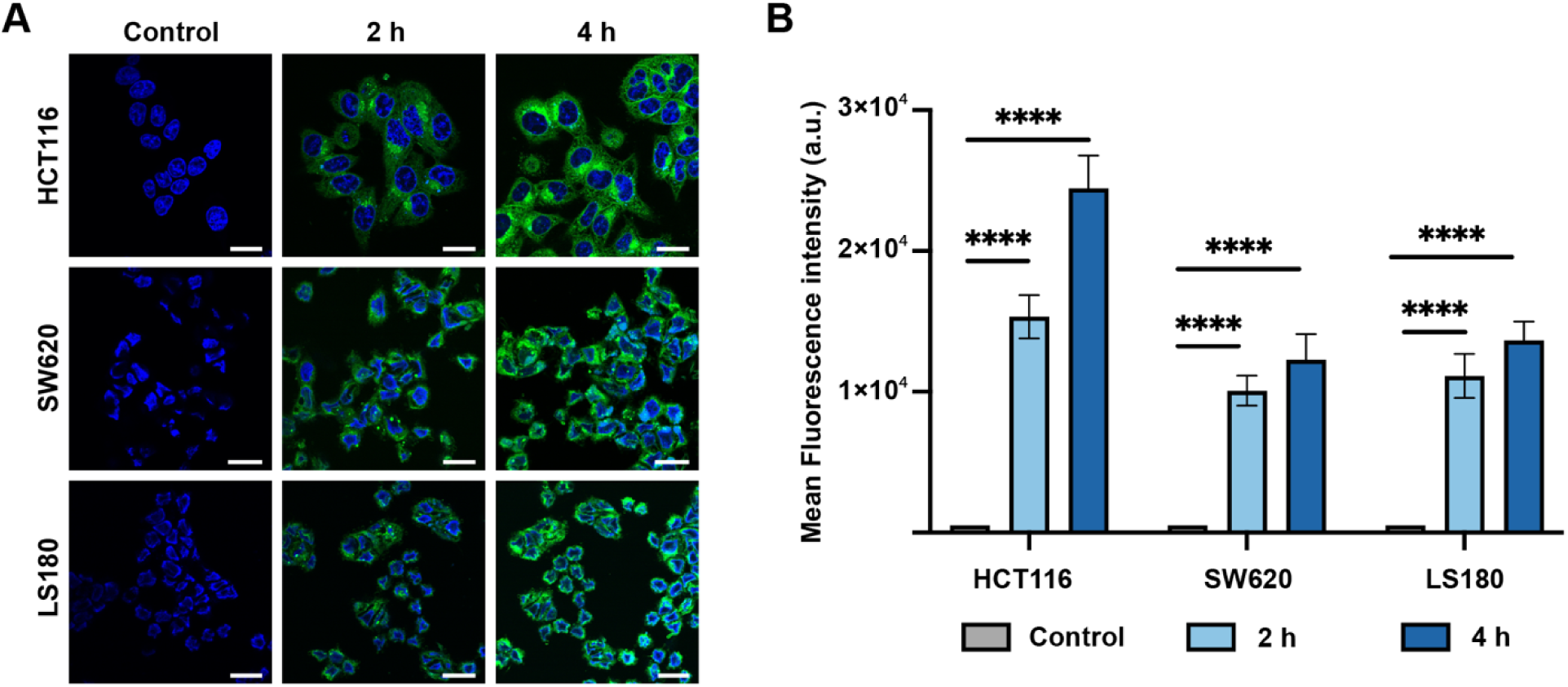
Cellular uptake of Cou6-labelled LNPs. (A) Representative confocal microscopy images of HCT116, SW620 and LS180 cells following incubation with Cou6-LNPs for 2 or 4 h. Untreated cells were included as controls. Nuclei were stained with DAPI (blue), and Cou6 fluorescence is shown in green. (B) Cellular uptake quantified as corrected total cell fluorescence (CTCF) per cell. Data are presented as mean ± SD from n = 3 independent experiments. Statistical comparisons used one-way ANOVA with Tukey’s multiple comparison test; ****P < 0.0001.

Quantitative corrected total cell fluorescence (CTCF) analysis further demonstrated increased intracellular fluorescence over time (Figure 2B). In HCT116 cells, CTCF increased from 15,327± 1,546 at 2 h to 24,431 ± 2,341 at 4 h. Corresponding increases were observed in SW620 cells (10,065 ± 1,065 to 12,256 ± 1,837) and LS180 cells (11,117 ± 1,562 to 13,639 ± 1,339), indicating differences in LNP uptake among the three cell lines. Overall, the optimised LNP formulation showed higher cellular uptake at 4 h, which was therefore selected for subsequent cellular evaluation

### *In vitro* KRAS gene silencing via siKRAS-LNPs

The gene silencing activity of siKRAS-LNPs was evaluated in KRAS-mutant HCT116 (G13D), SW620 (G12V) and LS180 (G12D) cells, as well as KRAS wild-type HT29 cells. RT-qPCR analysis showed that siKRAS-LNP treatment significantly reduced KRAS mRNA expression to 0.44-fold in HCT116, 0.34-fold in SW620 and 0.70-fold in LS180 cells compared with untreated controls (P < 0.001; Figure 3A-C). In contrast, siKRAS-LNP treatment did not reduce KRAS mRNA expression in HT29 cells (Figure 3D), demonstrating that the siKRAS exhibits mutation-selective silencing activity. Western blot analysis further confirmed reduced KRAS protein expression following siKRAS-LNP treatment in the three KRAS-mutant cell lines (Figure 3E-G).

**Figure 3.**
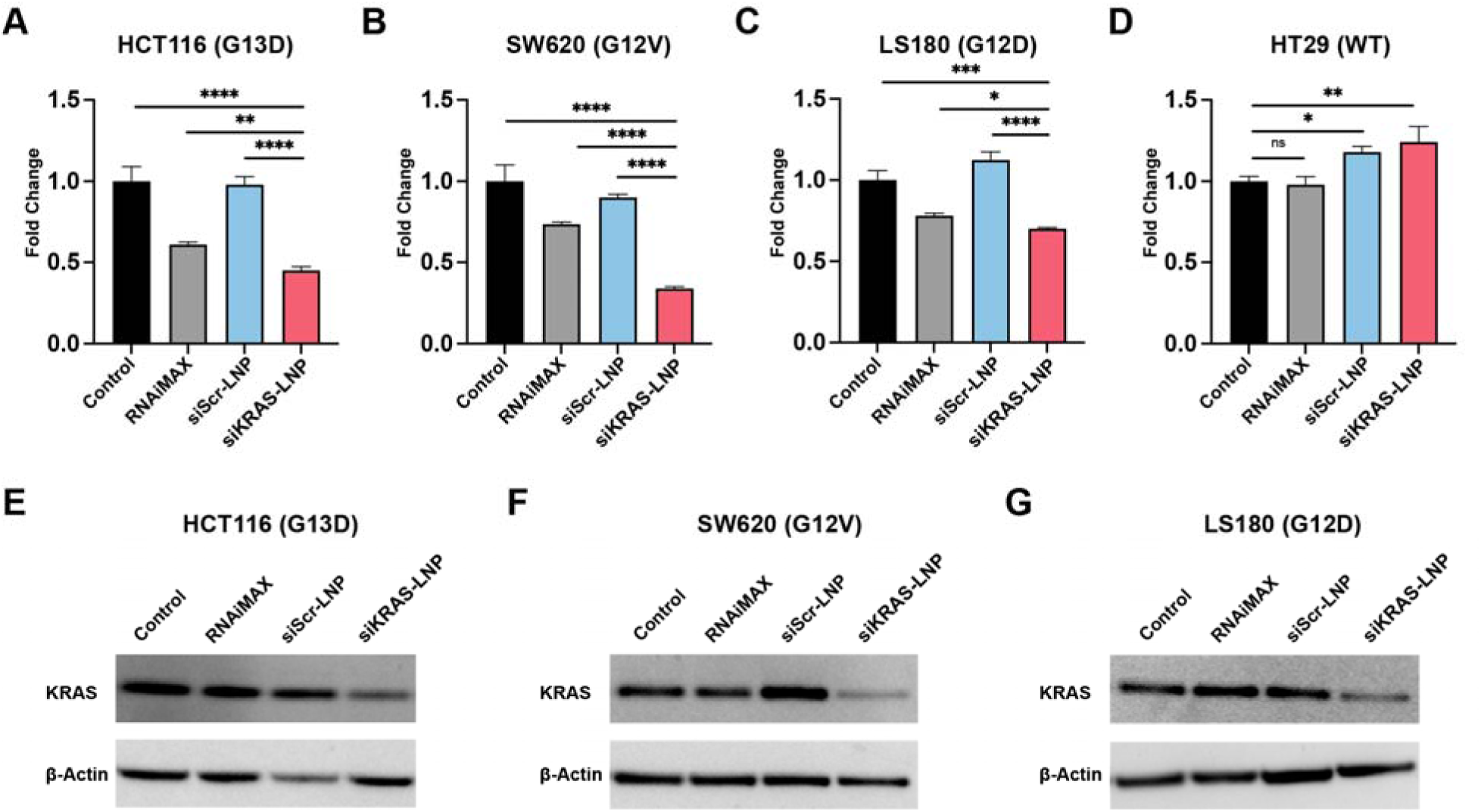
Evaluation of siKRAS-LNP-mediated gene silencing efficiency. (A-D) KRAS mRNA expression in HCT116, SW620, LS180, and HT29 cells following treatment with RNAiMAX, siScr-LNP, or siKRAS-LNP, normalized to untreated control. Data are presented as mean ± SD from n = 3 independent experiments. Statistical comparisons used one-way ANOVA with Tukey’s test; *P < 0.05, **P < 0.01, ***P < 0.001, ****P < 0.0001. (E-G) Representative Western blot images of KRAS in HCT116, SW620 and LS180 cells under the same treatment conditions. β-Actin was used as a loading control.

Importantly, siKRAS-LNPs achieved significantly higher KRAS silencing at both the mRNA and protein levels than RNAiMAX-mediated siKRAS transfection across all three KRAS-mutant cell lines. This enhanced silencing may reflect differences in siRNA delivery and intracellular availability between the two delivery systems, although the underlying mechanism was not directly investigated in this study.

### Cellular response to KRAS silencing

After confirming the silencing activity of siKRAS-LNPs *in vitro*, we next evaluated the functional effects of KRAS silencing on cellular behaviour. Cell proliferation was significantly reduced following siKRAS-LNP treatment compared with the other conditions, with the inhibitory effect increasing over time and being most pronounced in HCT116 cells (Figure 4A-C). Consistent with these results, siKRAS-LNP treatment reduced colony number and size after14 days (Figure 4D). Quantification of the surviving fraction confirmed reduced clonogenic survival in all three cell lines, with the greatest reduction observed in HCT116 cells, followed by LS180 and SW620 (Figure 4E-G). In contrast, siScr-LNP treatment had no significant effect. Cell migration was also reduced following siKRAS-LNP treatment, with relative migration rate reduced to 43.1 ± 1.4%, 57.1 ± 1.5% and 44.0 ± 6.3% in HCT116, SW620 and LS180, respectively (Figure 4H-I). Taken together, these results confirm that siRNA-mediated KRAS silencing reduces proliferation, colony formation, and migration of KRAS-mutant cells.

**Figure 4.**
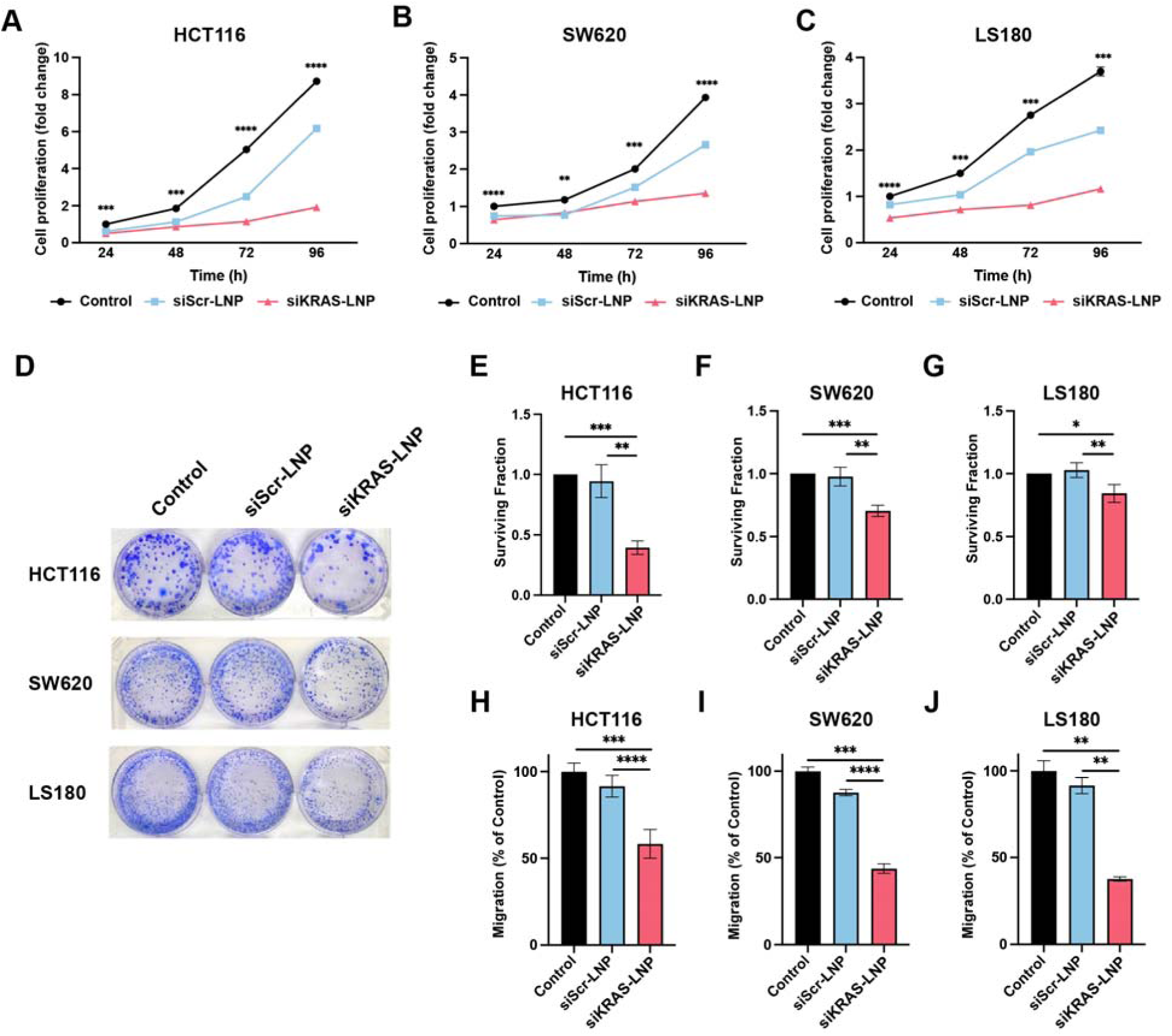
Assessment of cellular response following KRAS silencing. (A-C) Cell proliferation analysis of HCT116, SW620 and LS180 cells treated with siKRAS-LNP, siScr-LNP or untreated control over 96 h. (D) Representative colony formation images. (E-G) Clonogenic surviving fraction normalized to control. (H-J) Relative cell migration rate normalized to untreated control. Data are presented as mean ± SD from n = 3 independent experiments. Statistics: one-way ANOVA with Tukey’s test for (A-C); one-sample and two-sample t-tests for (E-J). *P < 0.05, **P < 0.01, ***P < 0.001, ****P < 0.0001.

### Transcriptomic profiling and pathway enrichment

To investigate the transcriptional response to KRAS silencing and identify associated downstream pathways, bulk RNA sequencing was performed on HCT116 cells treated with empty-LNP, siScr-LNP or siKRAS-LNP conditions. Principal component analysis showed clear clustering of biological replicates, while dispersion analysis showed consistency with the fitted mean-dispersion trend (Figure S1 and S2). Comparison of untreated and empty-LNP treated cells identified no differentially expressed genes, indicating minimal effects of the LNP carrier on the cellular transcriptome (Figure 5A). The subsequent analysis therefore focused on the siKRAS-LNP versus siScr-LNP comparison to identify transcriptional changes associated with KRAS silencing. This analysis identified 883 differentially expressed genes, including 495 upregulated and 388 downregulated genes (Figure 5B-C). Similar expression patterns were observed when siKRAS-LNP was compared with empty-LNP controls, indicating that the observed transcriptional changes were primarily associated with KRAS silencing rather than the LNP carrier (Figure S5).

**Figure 5.**
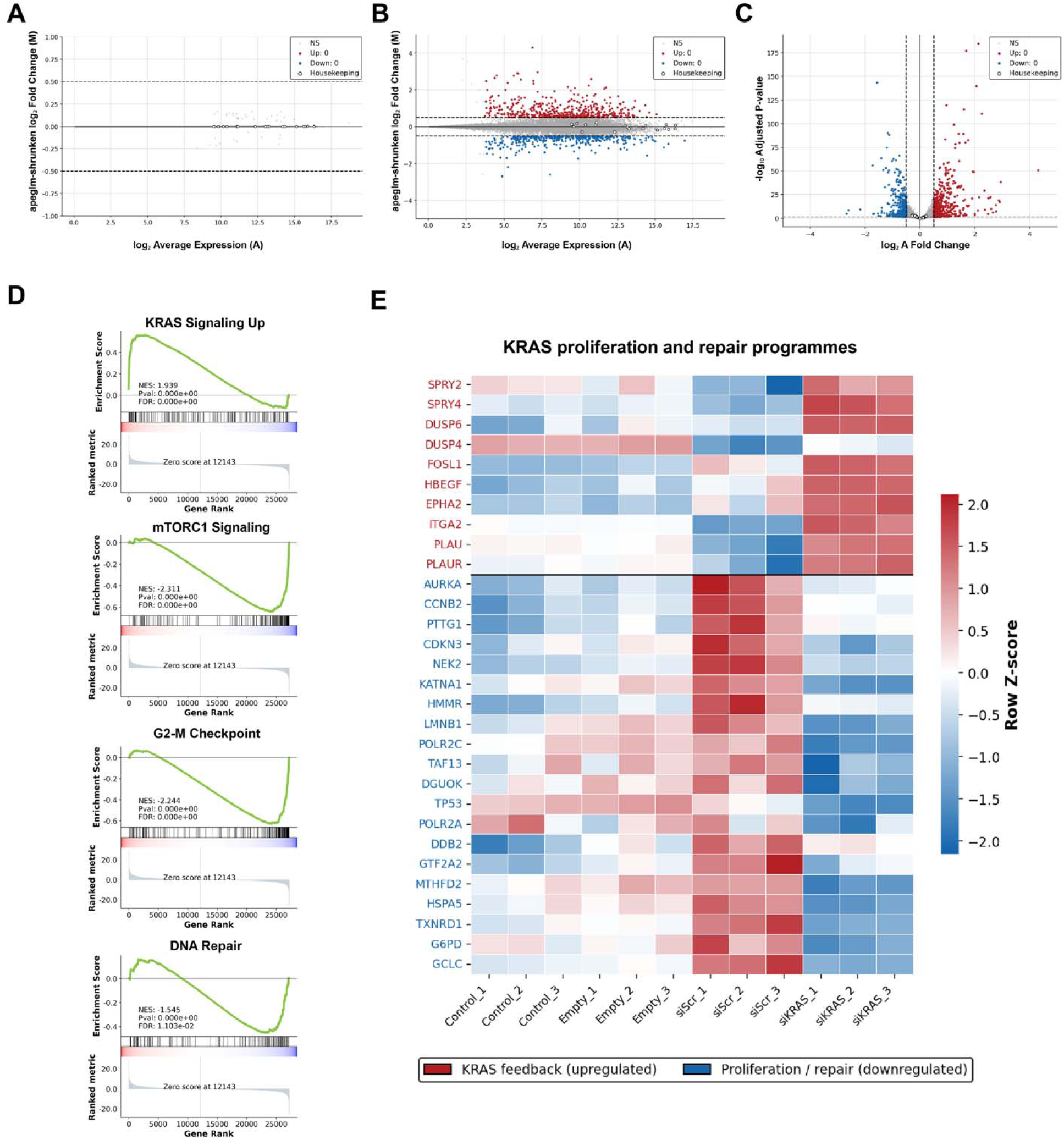
Evaluation of gene expression and pathway changes following KRAS silencing. (A) MA plot comparing untreated and empty-LNP cells. (B) MA and (C) volcano plots comparing siKRAS-LNP with siScr-LNP; differentially expressed genes were defined as baseMean ≥ 10, adjusted P < 0.05, and |log2 fold change| > 0.5. (D) Hallmark GSEA of mTORC1 signalling, G2/M checkpoint, DNA repair, and KRAS Signaling Up; NES and FDR values are shown. (E) Heatmap of representative KRAS pathway feedback and proliferation/repair genes, expressed as row Z-scores. RNA-seq: n = 3 biological replicates; DESeq2 Wald test with Benjamini– Hochberg correction. GSEA used 1,000 permutations.

Gene set enrichment analysis (GSEA) of siKRAS-LNP versus siScr-LNP showed negative enrichment of mTORC1 signalling (NES = −2.31, FDR < 0.001), G2/M checkpoint (NES = −2.24, FDR < 0.001) and DNA repair (NES = −1.55, FDR = 0.011) (Figure 5D). These changes were consistent with reduced activity of pathways associated with cell proliferation, metabolism, cell cycle and DNA repair. In contrast, KRAS Signaling Up gene set was positively enriched (NES = 1.94, FDR < 0.001), potentially reflecting an adaptive transcriptional feedback response following KRAS suppression (Figure 5D). The broader Hallmark enrichment profile is shown in Figure S4. To examine the gene-level changes underlying these enrichment patterns, representative genes were visualised by heatmap (Figure 5E). siKRAS-LNP treatment increased the expression of KRAS pathway feedback-associated genes, including SPRY2, SPRY4, DUSP6 and DUSP4, and reduced the expression of proliferation- and DNA repair-associated genes, including AURKA, CCNB2, HMMR, DDB2 and MTHFD2 (Figure 5E). Overall, these gene-level changes were consistent with the GSEA results (Figure 5D), supporting suppression of proliferative and DNA repair programmes together with increased expression of genes associated with KRAS pathway feedback.

Among the suppressed pathways, negative enrichment of DNA repair was of particular interest because of its potential therapeutic implications. Sub-pathway enrichment analysis using Reactome gene sets showed repression of multiple DNA repair processes, including base excision repair, nucleotide excision repair, non-homologous end joining, homologous recombination and DNA replication (Figure S6). These findings suggested that KRAS silencing may impair cellular responses to DNA damage and could therefore influence sensitivity to DNA-damaging chemotherapy. We therefore examined the relationship between the siKRAS transcriptional response and 5-fluorouracil (5-FU), a cornerstone chemotherapeutic agent for colorectal cancer [20]. 5-FU exerts its cytotoxic effects primarily through inhibition of thymidylate synthase, leading to dTTP depletion, uracil misincorporation into DNA and subsequent DNA damage [20]. The LINCS L1000 Fluorouracil Up gene set was enriched among siKRAS-upregulated genes (NES = 1.33, FDR = 0.023), whereas the Fluorouracil Down gene set was enriched among siKRAS-downregulated genes (NES = −1.42, FDR = 0.002) (Figure S7). These findings indicated that siKRAS treatment induced a transcriptional response overlapping with that associated with 5-FU exposure. Together with the suppression of DNA repair pathways, these findings provided a rationale for evaluating whether siKRAS-LNPs could enhance the response to 5-FU. This rationale motivated the subsequent combination studies.

### Cellular response to combined treatment of siKRAS-LNP and 5-fluorouracil

Building on the transcriptomic findings that KRAS silencing suppressed cell cycle and DNA repair pathways and showed overlap with 5-FU response signatures, we next investigated whether combining siKRAS-LNP with 5-FU could enhance treatment response in KRAS mutant CRC cells. As shown in Figure 6A-C, siKRAS-LNPs exhibited positive synergy with 5-FU across HCT116, SW620 and LS180 cells, with mean ZIP synergy scores of 14.26, 20.99, and 15.14, respectively. These positive ZIP scores indicate that the combination produced greater-than-additive effects across all three KRAS-mutant CRC cell lines, with the strongest synergy observed in SW620 cells. This finding supports the potential of combining KRAS silencing with 5-FU to enhance treatment response across prevalent KRAS-mutant CRC subtypes. Similar synergy patterns were observed using the HSA, Loewe and Bliss reference models (Figure S10).

**Figure 6.**
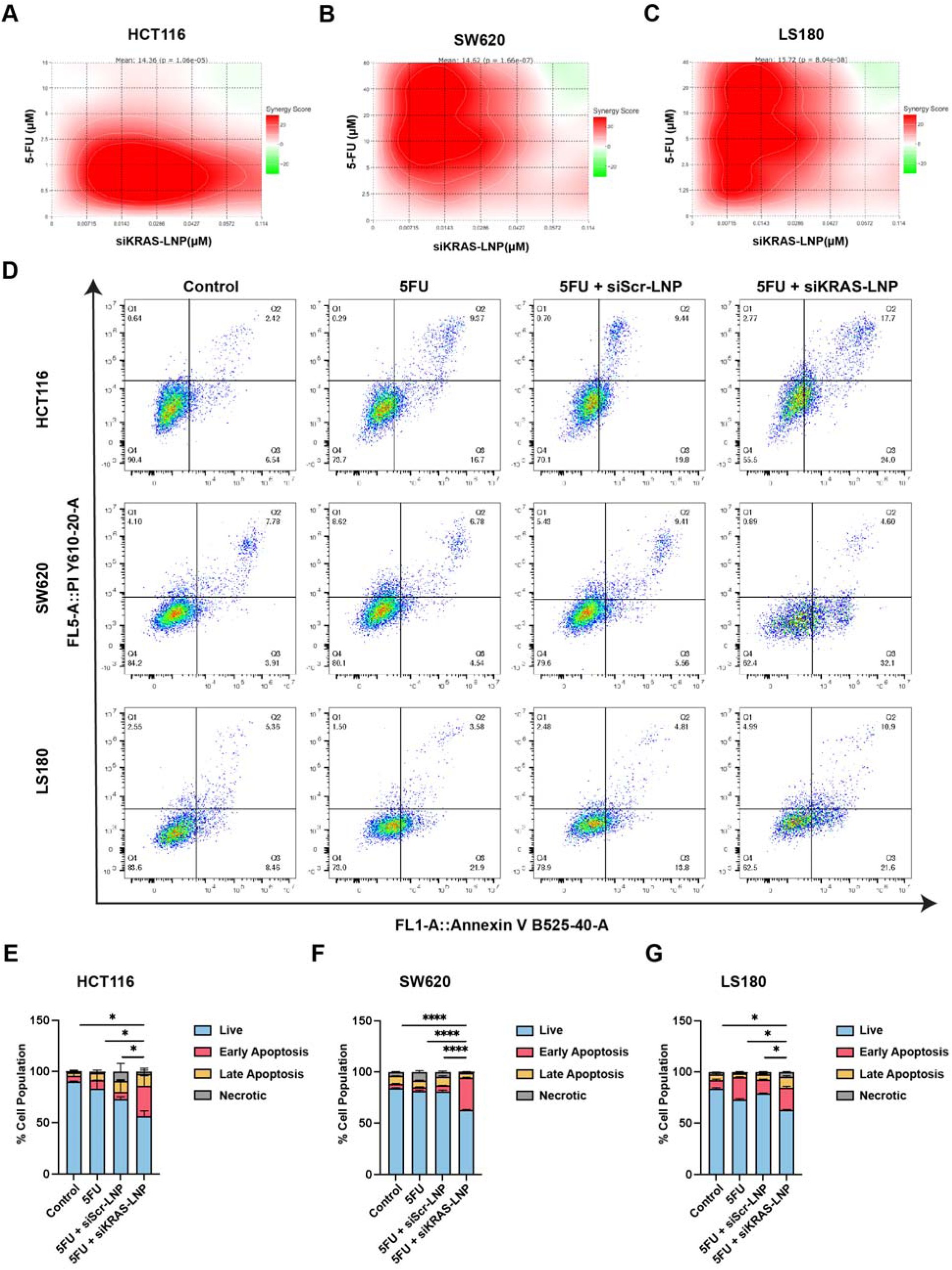
Synergy analysis and apoptosis following combined siKRAS-LNP and 5-fluorouracil treatment. (A-C) ZIP synergy maps for siKRAS-LNP and 5-FU in HCT116, SW620 and LS180 cells. Mean synergy scores are shown; scores >10 indicate synergy. (D) Annexin V/PI apoptosis analysis following treatment with 5-FU, 5-FU + siScr-LNP, or 5-FU + siKRAS-LNP. Representative dot plots (left) and quantification of live, early apoptotic, late apoptotic and necrotic populations (stacked bars). Data are presented as mean ± SD from n = 3 independent experiments. Apoptotic fractions were compared one-way ANOVA with Tukey’s multiple-comparisons test. *P < 0.05, **P < 0.01, ****P < 0.0001; ns, not significant.

Based on the dose response and synergy analysis, 28.6nM siKRAS-LNP was combined with the cell line specific IC concentration of 5-FU (HCT116, 1.97µM; SW620, 12.66µM; LS180, 7.8µM) for subsequent apoptosis analysis. Annexin V/PI staining showed that combined siKRAS-LNP and 5-FU treatment increased the apoptotic population to 41.3% ± 6.3% in HCT116, 36.6% ± 0.4% in SW620, and 32.4% ± 1.7% in LS180 (P = 0.013, P< 0.001 and P = 0.032; Figure 6D). These findings indicate that the combination of siKRAS-LNP and 5-FU significantly enhanced apoptosis compared with 5-FU treatment alone in all these three cell lines.

We also compared the siKRAS-LNP-induced transcriptional profile with a public dataset of 5-FU-resistant versus 5-FU-sensitive HCT116 cells (GSE153412; n = 3 per group) to examine whether KRAS silencing reversed the 5-FU resistance-associated transcriptional profile. No inverse correlation was observed across all genes or significantly altered genes (Figures S8 and S9), indicating that the enhanced apoptotic response was not associated with reversal of the established 5-FU resistance signature. Instead, it may be associated with the altered cell-cycle and DNA-repair states identified following KRAS silencing.

### Evaluation of combination treatment response in 3D spheroid

Having observed an enhanced apoptotic response to the combination treatment in monolayer cultures, we next assessed its anti-cancer effect in a 3D tumour model. We first examined the distribution of the developed LNPs in HCT116 spheroids using Cou6-LNPs. Green fluorescence was detected within the spheroids at both 24 and 48 h (Figure 7A), with mean fluorescence intensity increasing from 650.7 ± 19.0 a.u. at 24 h to 826.4 ± 23.3 a.u. at 48 h (Figure 7B), indicating time-dependent accumulation of LNPs within the spheroids. As substantial LNP accumulation was already evident at 24 h, this time point was selected for subsequent treatment studies.

**Figure 7.**
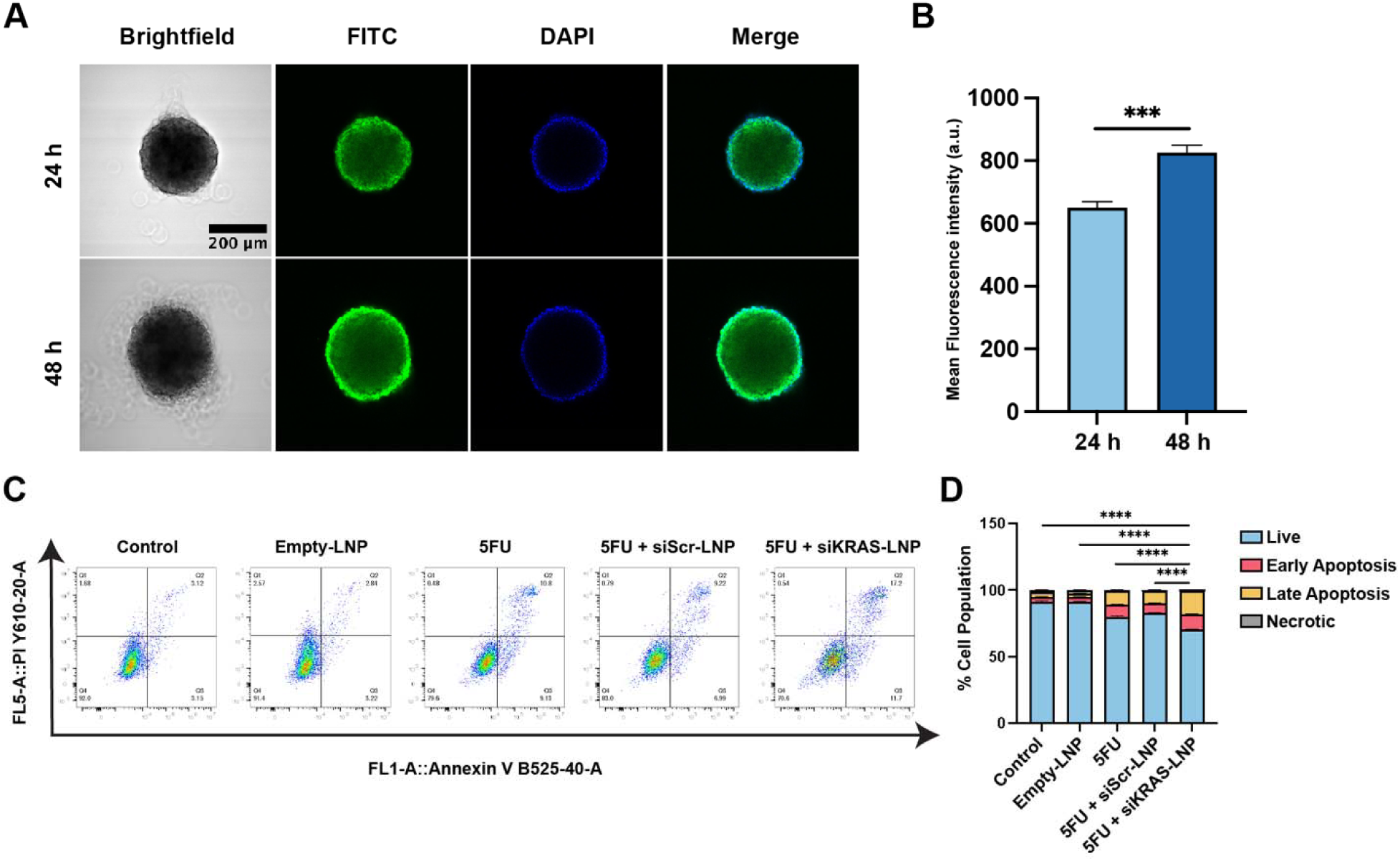
LNP penetration and treatment response in HCT116 spheroids. (A) Representative confocal images of HCT116 spheroids following incubation with Cou6-LNPs for 24 and 48 h. Nuclei are shown in blue and LNP associated fluorescence in green. Scale bar, 200 µm. (B) Mean fluorescence intensity per spheroid at 24 and 48 h.(C) Representative Annexin V/PI flow cytometry plots of dissociated HCT116 spheroids following treatments indicated in the figure. (D) Quantification of live, early apoptotic, late apoptotic and necrotic cell populations. Data are presented as mean ± SD from n = 3 independent experiments. Fluorescence intensity was compared using a two-sample t-test. Total apoptotic fractions were compared by one-way ANOVA with Tukey’s multiple-comparisons test; *P < 0.05, **P < 0.01, ***P < 0.001, ****P < 0.0001; ns, not significant.

The therapeutic response was subsequently evaluated by Annexin V/PI flow cytometry following treatments with empty-LNPs, 5-FU, 5-FU + siScr-LNPs or 5-FU + siKRAS-LNPs. As shown in Figure 7 C-D, the 5-FU + siKRAS-LNP method significantly increased the apoptotic population compared with all other treatment groups (6.2% vs 19.5% vs 16.3% vs 28.8%). These findings confirm an enhanced apoptotic response to the combination treatment method in the 3D spheroid model, further supporting its evaluation in *in vivo* studies.

## Discussion

KRAS remains a major therapeutic challenge in CRC, as current allele-selective inhibitors have primarily focused on the infrequent G12C, leaving other prevalent variants, including G12D, G12V and G13D, without established targeted therapies. In this study, we developed an siRNA-based therapeutic approach to suppress these three prevalent KRAS variants in CRC. The siKRAS-LNPs were formulated using DOTAP and the helper lipid DOPE, a lipid combination with an established history in nucleic acid delivery and prior clinical evaluation[21, 22].

Following siRNA loading, the LNPs exhibited physicochemical characteristics suitable for nucleic acid delivery, including a mean diameter of approximately 110 nm and an encapsulation efficiency of approximately 93%. The positive surface charge was consistent with the potential for interaction with negatively charged cell membranes and cellular delivery of the siRNA cargo [23, 24].

Previous KRAS RNAi studies have largely focused on pancreatic cancer models or individual KRAS variants, with more recent approaches mainly targeting G12V[18, 25, 26]. In contrast, our study demonstrates that a single siRNA sequence can suppress multiple KRAS variants in CRC cells. siKRAS-LNPs produced obvious KRAS silencing effect at both the mRNA and protein levels across three KRAS-mutant cell lines (Figure 3A-G). Importantly, the extent of KRAS silencing did not directly predict the functional response across these cell lines. Despite exhibiting the greatest KRAS knockdown, SW620 cells showed a weaker reduction in clonogenic survival and migration than HCT116 and LS180 cells (Figure 4E-J). This differential response may reflect variation in KRAS dependency across tumour backgrounds and KRAS alleles and reliance on alternative mitogen-activated signalling cascades in the face of KRAS inhibition [27, 28]. Differences in LNP uptake may also contribute, as the highest fluorescence level of LNPs was observed in HCT116 cells compared with SW620 and LS180 cells at the same time point (Figure 2B).

Transcriptomic profiling provided further insight into the molecular response to KRAS silencing. The transcriptional changes were characterised by suppression of proliferative and DNA-repair programmes, consistent with the reduced proliferation and clonogenic survival observed (Figure 4). In contrast, the positive enrichment of the KRAS Signaling Up gene set may reflect a residual KRAS signalling following partial silencing and a compensatory transcriptional response following KRAS suppression, consistent with adaptive RAS/MAPK feedback previously reported following KRAS inhibition [29, 30]. The suppression of G2/M checkpoint and DNA-repair pathways following KRAS silencing further supported the rationale for combining siKRAS-LNPs with 5-FU to enhance therapeutic efficacy. This approach produced synergistic effects across all three KRAS-mutant cells, with consistent interactions observed using the ZIP, HSA, Loewe and Bliss reference models (Figure 6A-C; Figure S10) [31]. The enhanced cellular apoptosis in both monolayer cultures and the 3D spheroid model was observed through this method (Figures 6D and 7C-D), supporting further evaluation of its therapeutic efficacy *in vivo*.

The current study also identifies several areas for further development. The present LNP formulation was developed as a proof-of-concept delivery system and has limited tumour-targeting capability. Further optimisation of the formulation and incorporation of tumour-targeting strategies will therefore be required to improve tissue-selective delivery. Co-delivery of siKRAS and 5-FU within a single delivery system could also be explored to achieve coordinated delivery of the two therapeutic agents. *In vivo* studies will be an important next step to evaluate tissue-level delivery, KRAS silencing, therapeutic efficacy and treatment tolerability.

Overall, this study demonstrates that an siRNA-based strategy can suppress three prevalent KRAS variants in CRC cell lines and enhance the apoptosis response to 5-FU chemotherapy. This combination approach may offer a therapeutic strategy for KRAS-mutant CRC subtypes that currently lack approved KRAS-targeted therapies. Further optimisation of tumour-selective delivery and *in vivo* validation will be essential to establish its therapeutic efficacy and translational potential.

## Materials and methods

### Materials

1,2-Dioleoyl-3-trimethylammonium-propane (DOTAP; Cat. 890890P), 1,2-dioleoyl-*sn*-glycero-3-phosphoethanolamine (DOPE; Cat. 850725P), coumarin-6 (Cat. 442631), MISSION® siRNA Universal Negative Control (siScr; Cat. SIC001), 5-Fluorouracil (5-FU; Cat. F6627) and the QCM™ 24-Well Fluorometric Migration Assay (Cat. ECM509) were purchased from Sigma-Aldrich Pty Ltd (Sydney, Australia). All lipids were dissolved in ethanol (≥99.9%, molecular biology grade) and stored at −20 °C. The mutant-selective siRNA EFTX-D1 (Silencer™ Custom Designed siRNA, Cat. 4399667; hereafter si*KRAS*), Lipofectamine™ RNAiMAX Transfection Reagent (Cat. 13778075), Opti-MEM™ I Reduced Serum Medium (Cat. 31985070), Dulbecco’s Modified Eagle Medium (DMEM; Cat. 11965092), fetal bovine serum (FBS; Cat. A5670701), Minimum Essential Medium (MEM; Cat. 11095080), McCoy’s 5A (modified) Medium (Cat. 16600082), Trypsin (0.25%) (Cat. 15050057), TrypLE™ Express (Cat. 12604021), the RediPlate™ 96 RiboGreen™ RNA Quantitation Kit (Cat. R32700), DAPI (Cat. R37606), and Nunc™ Sphera™ 96-well round (U)-bottom ultra-low-attachment plates (Cat. 174925) were purchased from Thermo Fisher Scientific (NSW, Australia). The RNeasy® Mini Kit (Cat. 74104) was purchased from QIAGEN (Hilden, Germany), and the Luna® Universal One-Step RT-qPCR Kit (Cat. E3006S) from New England Biolabs (Ipswich, MA, USA).

For Western blotting, the Mini Gel Tank (Cat. A25977), Mini Blot Module (Cat. B1000), nitrocellulose membrane (Cat. LC2000), Pierce™ ECL Western Blotting Substrate (Cat. 32209), NuPAGE™ 4–12% Bis-Tris protein gels (Cat. NP0321BOX), NuPAGE™ Transfer Buffer (20×) (Cat. NP00061), NuPAGE™ MOPS SDS Running Buffer (20×) (Cat. NP0001), RIPA Lysis and Extraction Buffer (Cat. 89900), Halt™ Protease and Phosphatase Inhibitor Cocktail (Cat. 78440), and NuPAGE™ LDS Sample Buffer (4×) (Cat. NP0007) were purchased from Thermo Fisher Scientific (NSW, Australia). Tween® 20 (Cat. 3005), anti-KRAS antibody (Cat. 33197S), anti–β-actin antibody (Cat. 4967S) and anti-rabbit IgG, HRP-linked Antibody (Cat 7074) were purchased from Cell Signaling Technology (MA, USA). All other reagents were of analytical grade unless otherwise stated, and nuclease-free water was used throughout for RNA work.

### Cell culture

Human colorectal cancer cell lines HCT116, SW620, LS180, and HT29 were purchased from the American Type Culture Collection (ATCC, USA). HCT116 and HT29 cells were cultured in McCoy’s 5A (Modified) Medium, SW620 cells were cultured in Dulbecco’s Modified Eagle Medium (DMEM), and LS180 cells were cultured in Minimum Essential Medium. All culture media were supplemented with 10% fetal bovine serum (FBS) and 1% penicillium/streptomycin antibiotic. Cells were maintained at 37°C in a humidified incubator with 5% CO and were passaged upon reaching approximately 80–90% confluence using 0.25% trypsin-EDTA. Cells were tested for mycoplasma and only mycoplasma-free cells were used for experiments.

### Preparation of siRNA-loaded LNPs

LNPs were prepared by microfluidic mixing using NanoAssemblr Ignite system (Cytiva, USA). The organic phase consisted of DOTAP and DOPE, each dissolved in ethanol at 25 mg/mL and combined at 1:1 molar ratio. The aqueous phase consisted of 100 pmol of siRNA, either siKRAS or siScr, in 10 mM Tris buffer (pH 7.4). The aqueous and organic phases were mixed at a flow rate ratio of 1:1 and a total flow rate of 12 mL/min. Following mixing, LNPs were purified and buffer-exchanged by centrifugal ultrafiltration using 100 kDa molecular weight cut-off filter at 8,000 × g for 10 min to remove residual ethanol and unencapsulated siRNA. Fluorescent LNPs were prepared using the same protocol with coumarin-6 incorporated into the organic phase at 0.48 mol% of total lipid content; while empty LNPs were prepared with a siRNA-free aqueous phase. All purified LNPs were stored at 4 °C for subsequent experiments.

### Physicochemical characterisation of siRNA-loaded LNPs

The hydrodynamic diameter (Z-average), polydispersity index (PDI), and zeta potential were measured by Malvern Zetasizer (Malvern Panalytical Co., UK). Nanoparticle morphology was examined by cryo-electron microscopy (Cryo-TEM). The encapsulation efficiency of siRNA (EE%) was quantified using RiboGreen Assay (Thermo Fisher Scientific) according to the manufacturer’s instructions. Fluorescence was measured before and after disruption of the nanoparticles with 0.5% Triton X-100, and encapsulation efficiency was calculated as EE% = (total siRNA – free siRNA) / total siRNA × 100. To evaluate the stability of siRNA-LNP under different conditions, 100 μL of LNPs were diluted in 1 mL of serum-free culture medium, culture medium supplemented with 10% FBS, or buffers at pH 5, 6, and 7. Samples were incubated at 37°C, and the hydrodynamic diameter and PDI were measured at 0, 4 and 24 h using DLS.

### Cellular uptake of LNPs

Cellular uptake of LNPs was evaluated by confocal microscopy using coumarin-6 loaded LNPs. HCT116, SW620, and LS180 cells were seeded at a density of 2 × 10 cells per glass-bottom dish and cultured for 24 h at 37°C to allow cell attachment. Cells were then incubated with coumarin-6 loaded LNPs (2.92 µg/mL) in complete culture medium for 2h and 4h. Following incubation, cells were washed with Dulbecco’s phosphate-buffered saline (DPBS; 1×, pH 7.4) and stained with Hoechst 33342 for 10 min. Live-cell imaging was performed using a Nikon A1R laser scanning confocal microscope (Nikon Corporation, Japan) with a 60× oil immersion objective. Cellular uptake was quantified by measuring the corrected total cell fluorescence (CTCF) using Fiji software.

### KRAS gene silencing and protein analysis

For gene knockdown studies, HCT116, SW620, LS180 and HT29 cells were seeded at a density of 5 × 10 cells/well in 6-well plates and cultured for 24 h prior to treatment. Cells were treated with siKRAS-LNPs or siScr-LNPs in Opti-MEM Reduced Serum Medium for 4 h, after which the treatment medium was replaced with complete culture medium. Where indicated, Lipofectamine™ RNAiMAX complexed with the corresponding siRNAs was used as a positive transfection control. KRAS knockdown was assessed at the mRNA level after 24 h and at the protein level after 72 h.

Total RNA was extracted using the RNeasy Mini Kit according to the manufacturer’s instructions. RNA concentration and purity were determined using a NanoDrop™ One Microvolume UV-Vis Spectrophotometer (Thermo Fisher Scientific, Australia). KRAS mRNA expression was quantified using the Universal One-Step RT-qPCR Kit (New England Biolabs) on a QuantStudio™ 6 Pro Real-Time PCR System (Thermo Fisher Scientific, Australia). Gene expression was normalised to GAPDH using the 2^;−ΔΔCt method. All reactions were performed in technical triplicate using at least three independent biological replicates. Primer sequences (5′– 3′) were used as follows: KRAS forward, TGACCTGCTGTGTCGAGAAT; KRAS reverse, TTGTGGACGAATATGATCCAA; GAPDH forward, CGCTCTCTGCTCCTCCTGTTC; GAPDH reverse, ATCCGTTGACTCCGACCTTCAC.

For western blot analysis, cells were lysed in RIPA Lysis and Extraction Buffer (Thermo Fisher Scientific) supplemented with Halt™ Protease and Phosphatase Inhibitor Cocktail. Protein concentration was determined using the Pierce™ BCA Protein Assay Kit. Equal amounts of protein (20 μg) were mixed with NuPAGE™ LDS Sample Buffer (4×), loaded on NuPAGE™ 4– 12% Bis-Tris gels in NuPAGE™ MOPS SDS Running Buffer using a Mini Gel Tank and transferred onto nitrocellulose membranes using NuPAGE™ Transfer Buffer and a Mini Blot Module. Membranes were blocked with 5% bovine serum albumin (BSA) in Tris-buffered saline containing 0.1% Tween® 20 (TBST) and incubated with primary antibodies against KRAS protein (1:1000), followed by an HRP-conjugated anti-rabbit IgG secondary antibody (1:5000). Protein bands were visualized using Pierce™ ECL Western Blotting Substrate and imaged with an Amersham ImageQuant™ 800 (Cytiva, Marlborough, MA, USA). Band intensities were quantified using Fiji software and normalized to β-actin.

### Transcriptomic analysis

HCT116 cells were treated with siKRAS-LNPs, siScr-LNPs or empty-LNPs for 24 h, and total RNA was extracted using the RNeasy Mini Kit (QIAGEN). RNA samples were submitted to Novogene for bulk RNA-sequencing, and subsequent bioinformatic and statistical analyses were performed in-house. Unstranded, poly(A)-selected RNA-sequencing libraries were prepared and sequenced on an Illumina platform using 150-bp paired-end reads. Read quality was assessed using FastQC (v0.12.1) and summarised using MultiQC. Reads were aligned to the human reference genome GRCh38.p14 using STAR (v2.7.11b), with the NCBI RefSeq annotation and the ––quantMode GeneCounts option. Gene-level read counts were generated using an unstranded counting strategy.

Differential expression analysis was performed using DESeq2 (PyDESeq2 v0.5.2). Genes with zero counts across all samples were excluded before analysis, with no additional low-expression filtering applied. Statistical significance was assessed using the Wald test, with P values adjusted for multiple testing using the Benjamini–Hochberg method. Log2 fold changes were shrunk using the adaptive apeglm prior with the lfcShrink function. Genes with a baseMean ≥ 10, adjusted P < 0.05, and log2 fold change > 0.5 were defined as differentially expressed.

GSEA was performed using the prerank implementation in GSEApy (v1.2.1). Genes were ranked by the DESeq2 Wald statistic and analysed against MSigDB Hallmark, Reactome, and LINCS L1000 gene sets using 1,000 permutations (seed = 6). DNA repair pathways were analysed using Reactome gene sets, and 5-FU response signatures using LINCS L1000 Fluorouracil Up and Down signatures. Over-representation analysis was performed using Enrichr against KEGG, MSigDB Hallmark, MSigDB Oncogenic Signatures, Reactome, WikiPathways and BioCarta, with adjusted P < 0.05 as the significance threshold. Local GMT files were used for reproducibility.

### *In vitro* functional assays

Cell viability was evaluated using the MTT Assay (Thermo Fisher Scientific, Cat. 475989). HCT116, SW620, and LS180 cells were seeded at 1 × 10 cells/well in 96-well plates and cultured for 24 h before treatment. Cells were incubated with different LNP samples for 4 h, after which the medium was removed and the cells were maintained in fresh complete culture medium for an additional 24 h. Cell viability was assessed after 24, 48, and 72 h by incubation with MTT reagent according to the manufacturer’s instructions, and absorbance was measured at 570 nm using a Tecan Infinite microplate reader (Tecan, Switzerland). Cell viability was normalized to untreated controls, and IC values were calculated by nonlinear regression using GraphPad Prism.

For colony formation assays, cells were seeded in 6-well plates and cultured for 24 h prior to treatment. Following a 4 h treatment period, the medium was replaced with fresh complete culture medium, and cells were incubated for an additional 24 h. Cells were then harvested and re-seeded at 500 cells/well in 6-well plates and cultured for 14 days, with medium refreshed twice weekly. Colonies were fixed with 10% formaldehyde for 30 min, stained with 0.1% crystal violet (Sigma-Aldrich), and colonies containing more than 50 cells were counted manually.

Cell migration was assessed using the QCM™ 24-Well Fluorometric Cell Migration Assay (Sigma-Aldrich, Cat. ECM509). Following treatment, cells were resuspended in serum-free culture medium and seeded at 5 × 10 cells/insert into the upper chamber. Medium supplemented with 10% FBS was added to the lower chamber as a chemoattractant. After 24 h, migrated cells were detached, labelled with the supplied CyQuant® GR dye, and quantified by fluorescence (Ex/Em = 480/520 nm) using a Tecan Infinite microplate reader (Tecan, Switzerland).

### Synergy and apoptosis analysis

Cells were first incubated with siKRAS-LNPs (28.6 nM) for 4 h, after which the treatment medium was replaced with complete culture medium. Following an additional 24 h incubation, cells were treated with a range of 5-FU concentrations (HCT116: 0, 0.5, 1, 2.5, 5, 10, 15 µM; SW620: 0, 2.5, 5, 10, 20, 40, 80 µM; LS180: 0, 1.25, 2.5, 5, 10, 20, 40 µM) for 72 h. Cell viability was determined using the MTT assay, and drug interactions were analysed using SynergyFinder based on the Highest Single Agent (HSA), Bliss Independence, Loewe Additivity and Zero Interaction Potency (ZIP) reference models. Synergy scores were visualized as two-dimensional heat maps, and the optimal drug combination identified from the synergy analysis was selected for subsequent apoptosis studies.

For apoptosis analysis, HCT116, SW620 and LS180 cells were seeded at 5 × 10 cells/well in 6-well plates and cultured for 24 h prior to treatment. Cells were treated with siKRAS-LNPs (28.6 nM) for 4 h before replacement with complete culture medium. After an additional 24 h, cells were treated with the IC concentration of 5-FU (HCT116, 1.97 µM; SW620, 12.66 µM; LS180, 7.8 µM) and incubated for 72 h. Apoptosis was quantified after 72 h using the BD Pharmingen™ FITC Annexin V Apoptosis Detection Kit I (BD Biosciences) according to the manufacturer’s instructions. Cells were stained with Annexin V-FITC and propidium iodide (PI) and analysed using a CytoFLEX LX Flow Cytometer (Beckman Coulter, USA). Data were analysed using FlowJo software (FlowJo 10.8.1).

### Three-dimensional tumour spheroid studies

Three-dimensional (3D) tumour spheroids were generated from HCT116 cells using Nunc™ Sphera™ 96-well round-bottom ultra-low attachment plates (Thermo Fisher Scientific, Cat. 174925). Cells were seeded at a density of 1 × 10³ cells/well in complete culture medium and cultured for 4 days to allow compact spheroid formation.

For nanoparticle penetration studies, spheroids were incubated with coumarin-6-labeled LNPs (2.92 µg/mL) for up to 48 h. At 24 and 48 h, spheroids were stained with Hoechst 33342 for 30 min, washed twice with DPBS (1×, pH 7.4) and imaged using a Nikon A1R laser-scanning confocal microscope (Nikon Corporation, Japan). Confocal Z-stack images were acquired to assess nanoparticle penetration throughout the spheroid, and fluorescence intensity was quantified using Fiji software.

For viability studies, spheroids were treated with the selected combination of siKRAS-LNPs and 5-FU and cultured for the indicated treatment period. Spheroids were subsequently dissociated into single cell suspensions, and apoptosis was quantified using the BD Pharmingen™ FITC Annexin V Apoptosis Detection Kit I (BD Biosciences, Cat. 556547) as described above. Samples were analysed using a CytoFLEX LX Flow Cytometer (Beckman Coulter, USA), and data were processed using FlowJo (version 10.8.1).

### Statistical analysis

All statistical analyses were performed using GraphPad Prism (version 10.2.0; GraphPad Software, USA). Data are presented as mean ± standard deviation (SD) from three independent biological replicates (n = 3), unless otherwise stated. Comparisons between two groups were performed using an unpaired two-tailed Student’s t-test, while comparisons among more than two groups were analysed using ordinary one-way analysis of variance (ANOVA) followed by Tukey’s multiple comparisons test. A P value < 0.05 was considered statistically significant.

## Data availability statement

All analysis code is available on GitHub at https://github.com/SS-Steve/siKRAS_5FU_CRC_RNAseq. Raw and processed RNA-seq data generated in this study will be deposited in the NCBI Gene Expression Omnibus (GEO) upon publication. All other data supporting the findings of this study are available within the article and its supplementary information or from the corresponding author upon reasonable request.

## Acknowledgments

The authors acknowledge the Microbial Imaging Facility at the University of Technology Sydney for access to confocal microscopy facilities and technical support with image acquisition.

## CRediT authorship contribution statement

Yuhang Wang: Writing – review & editing, Writing – original draft, Visualization, Validation, Methodology, Investigation, Formal analysis, Data curation. Shuquan Su: Writing – review & editing, Methodology. Gyorgy Hutvagner: Writing – review & editing. Kainat Zahra: Writing – review & editing. Rui Sang: Writing – review & editing. Mark Molloy: Writing – review & editing. Wei Deng: Conceptualization, Project administration, Methodology, Funding acquisition and Writing – review & editing.

