## Supplemental Information for "Lipid nanoparticle-mediated RNAi targeting of KRAS mutants sensitises colorectal cancer cells to chemotherapy"

**Table S1.** Table for LNP size and PDI formulations tested

| No. | DOTAP:DOPE Mole Ratio | Buffer | Flow Rate Ratio (FRR) | Total Flow Rate (TFR) | Purification | Z-Average (d.nm) | PDI |
| --- | --- | --- | --- | --- | --- | --- | --- |
| 1 | (1:1) | PBS | (1:1) | 12ml/min | dialysis | 517.6 | 0.13 |
| 2 | (1:1) | TRIS | (1:1) | 12ml/min | filter | 70.75 | 0.169 |
| 3 | (1:1) | TRIS | (3:1) | 12ml/min | filter | 1047 | 0.927 |
| 4 | (1:1) | TRIS | (5:1) | 12ml/min | filter | 394.7 | 0.578 |
| 5 | (1:1) | TRIS | (1:1) | 5ml/min | filter | 115.6 | 0.353 |
| 6 | (1:1) | TRIS | (1:1) | 20ml/min | filter | 230.3 | 0.419 |
| 7 | (1:1) | TRIS | (1:1) | 12ml/min | dialysis | 317.7 | 0.13 |
| 8 | (1:2) | TRIS | (1:1) | 12ml/min | dialysis | 681.5 | 0.018 |
| 9 | (2:1) | TRIS | (1:1) | 12ml/min | dialysis | 131.6 | 0.197 |
| 10 | (3:1) | TRIS | (1:1) | 12ml/min | dialysis | 82.56 | 0.173 |
| 11 | (4:1) | TRIS | (1:1) | 12ml/min | dialysis | 186.1 | 0.284 |

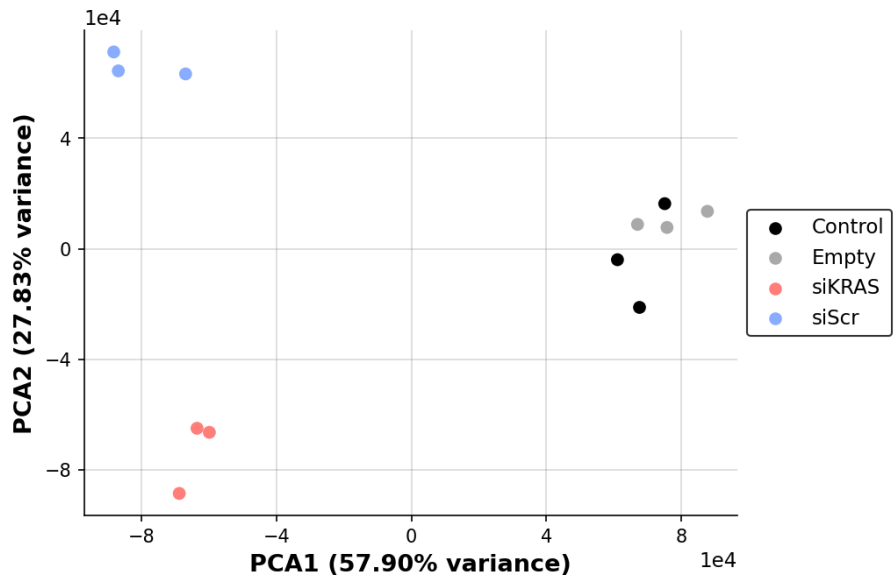

**Figure S1.** RNA-seq quality control. Principal component analysis of normalised expression across all 12 samples, coloured by treatment group, showing clear clustering of biological replicates.

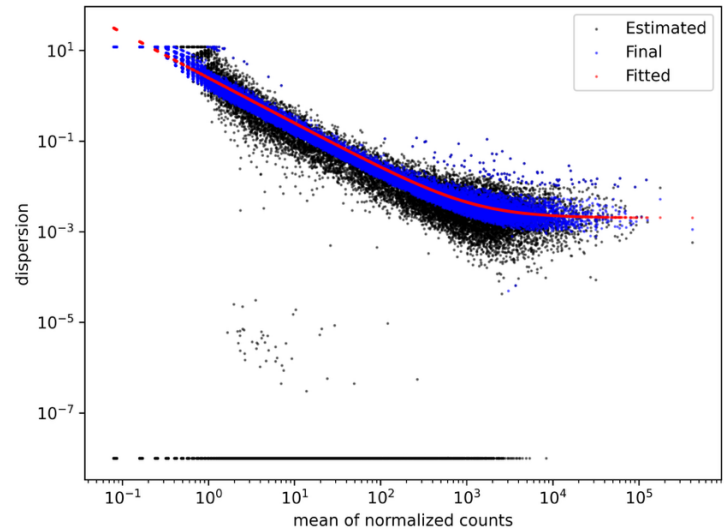

**Figure S2.** DESeq2 dispersion estimates. Plot of gene-wise dispersion against mean of normalised counts, showing the fitted dispersion trend, confirming appropriate mean–variance modelling of the negative binomial distribution.

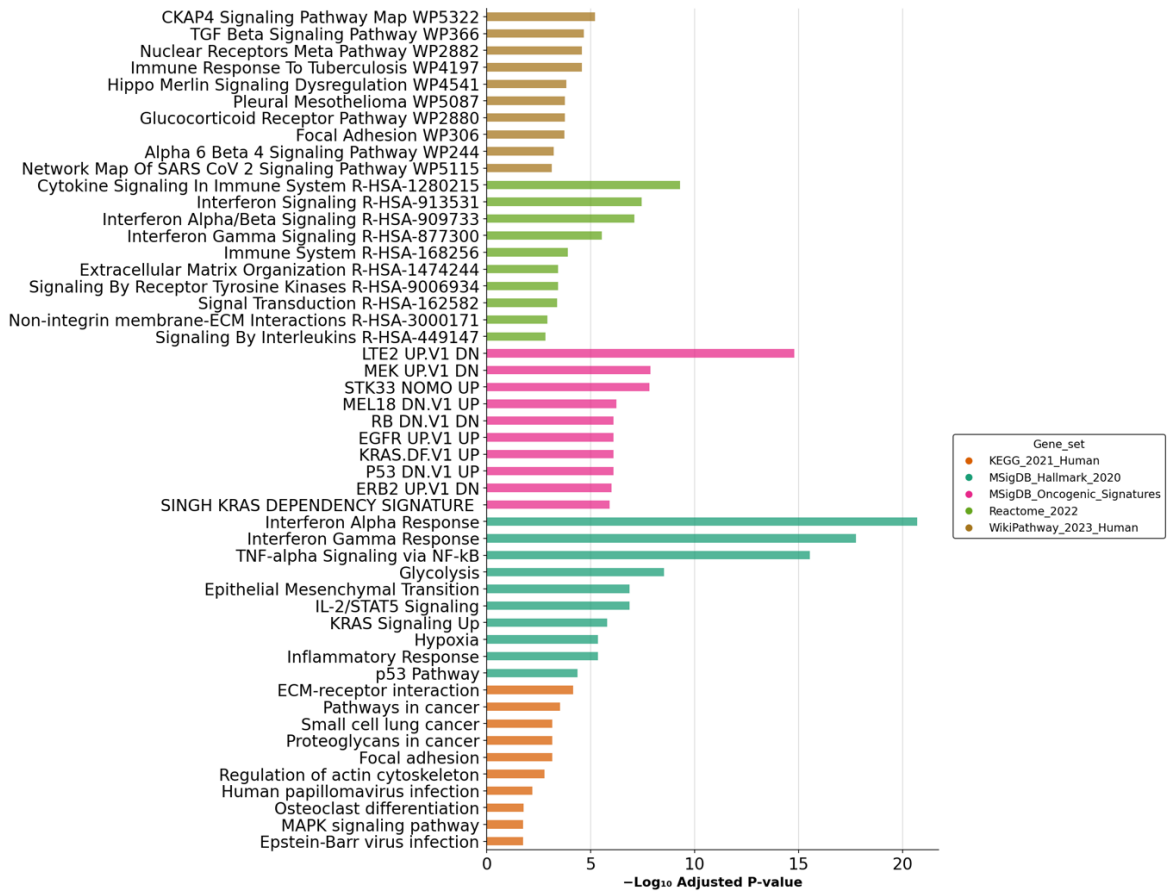

**Figure S3.** Over-representation analysis of differentially expressed genes using Enrichr across curated pathway databases (KEGG, MSigDB Hallmark, MSigDB Oncogenic Signatures, Reactome, WikiPathways, BioCarta), showing the top enriched terms ranked by adjusted P value.

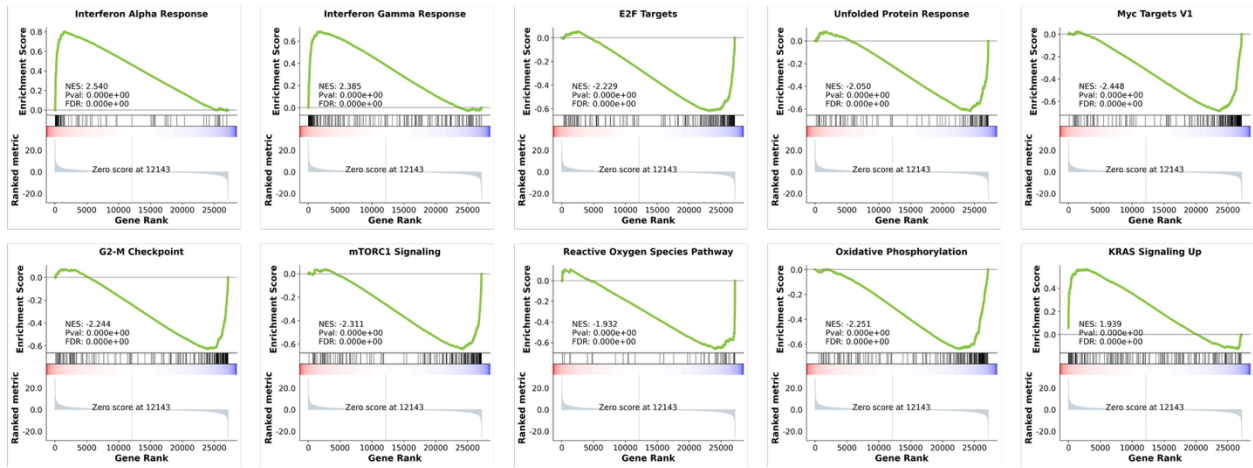

**Figure S4.** Complete GSEA results for the top 10 most significantly enriched Hallmark gene sets, providing the full enrichment landscape beyond the representative pathways shown in Figure 5D.

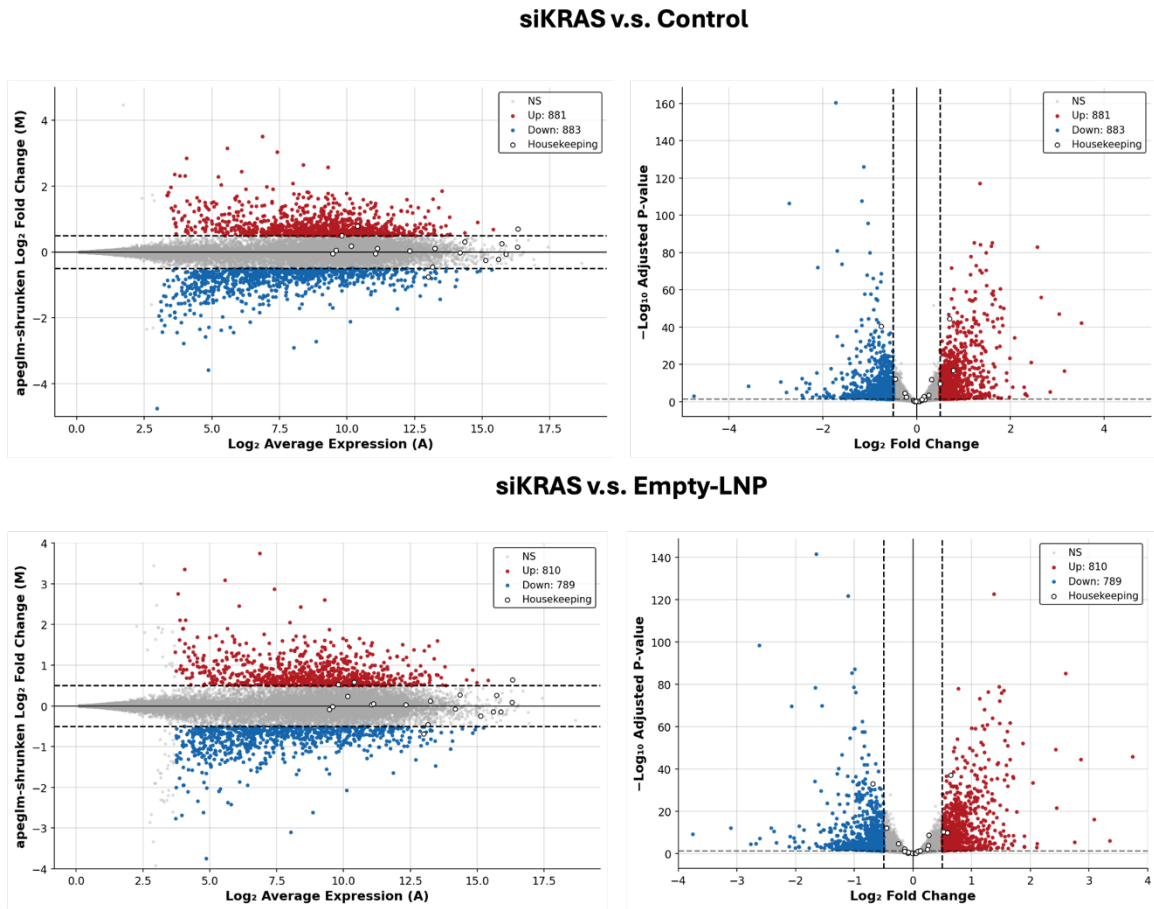

**Figure S5.** Differential expression for additional contrasts. MA plots and volcano plots for siKRAS versus Empty-LNP and siKRAS versus Control, showing that the transcriptomic changes are consistent across different reference conditions.

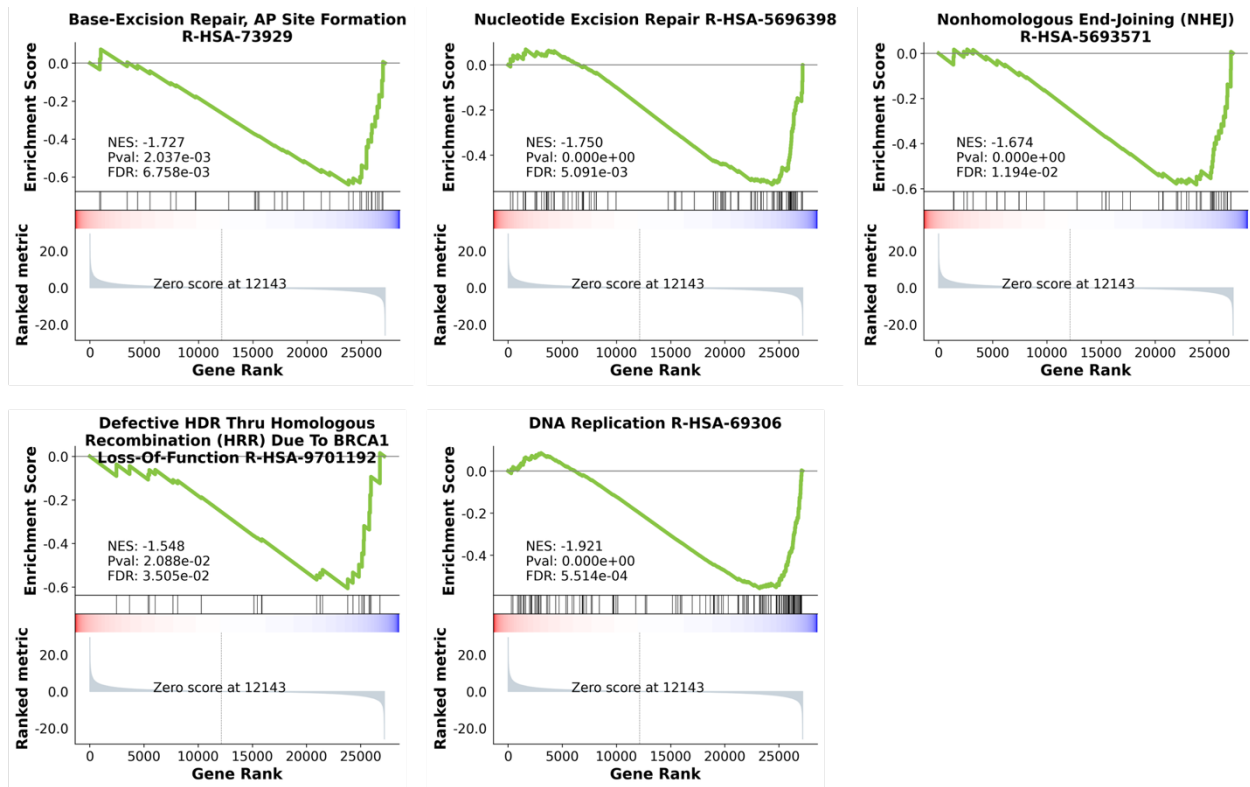

**Figure S6.** GSEA of DNA repair sub-pathways, revealing coordinate repression of base excision repair (AP site formation), nucleotide excision repair, non-homologous end joining, homologous recombination repair, and DNA replication.

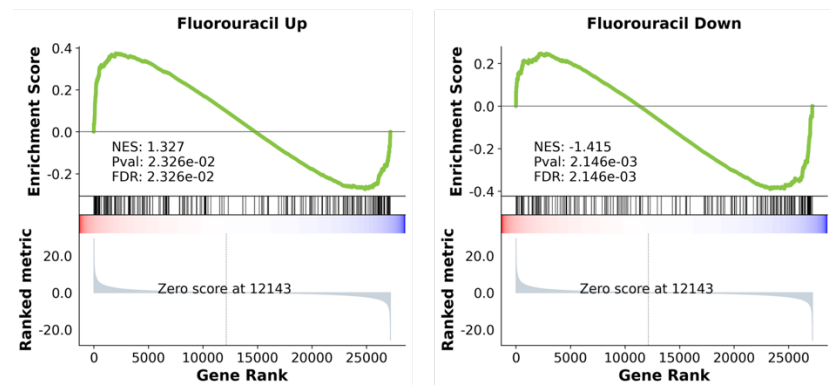

**Figure S7.** Overlap of the siKRAS transcriptome with the LINCS L1000 5-FU response signatures, showing enrichment of the Fluorouracil Up gene set among siKRAS-upregulated genes and the Fluorouracil Down gene set among siKRAS-downregulated genes.

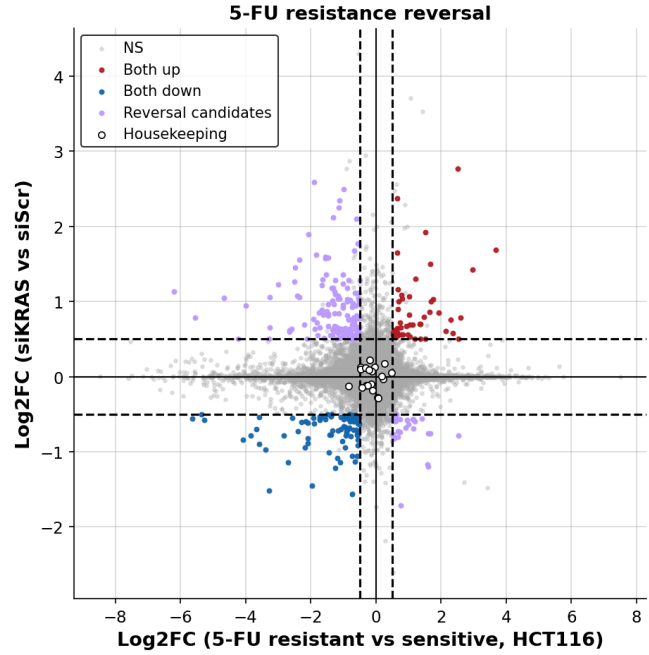

**Figure S8.** Scatter plot comparing 5-FU resistance log2 fold changes (GSE153412, resistant vs sensitive HCT116) with siKRAS log2 fold changes, coloured by concordance (Both up, Both down, Reversal candidates). Housekeeping genes are shown as white-filled circles.

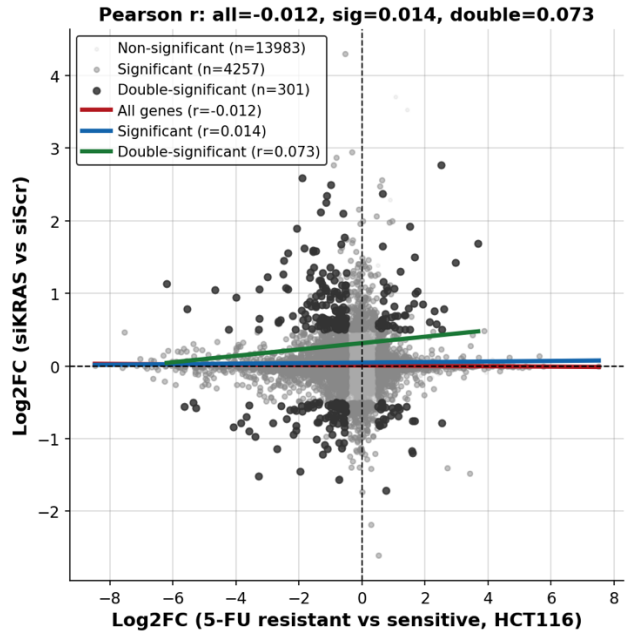

**Figure S9.** Pearson correlation between resistance and siKRAS log2 fold changes, computed for all genes, significant genes (union), and double-significant genes, with a linear fit line per group; all three correlations are near zero, indicating that siKRAS does not reverse the resistance signature.

### HCT116

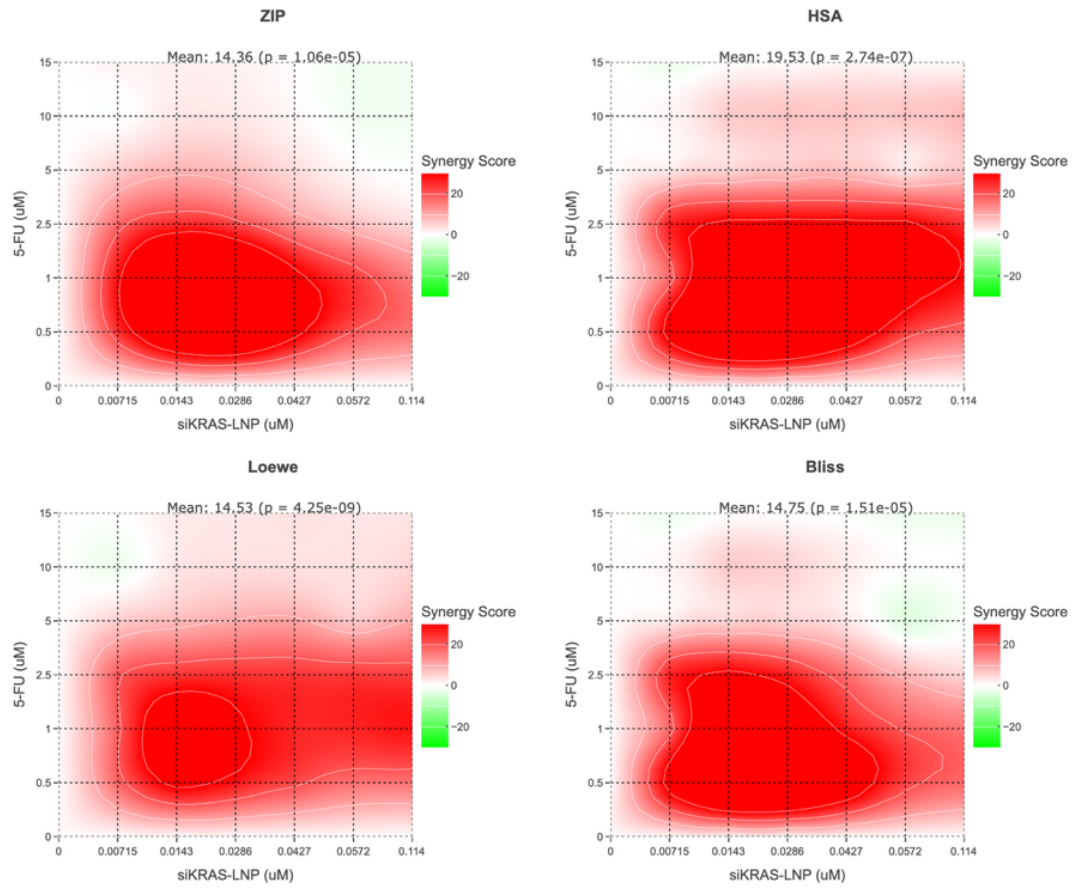

## SW620

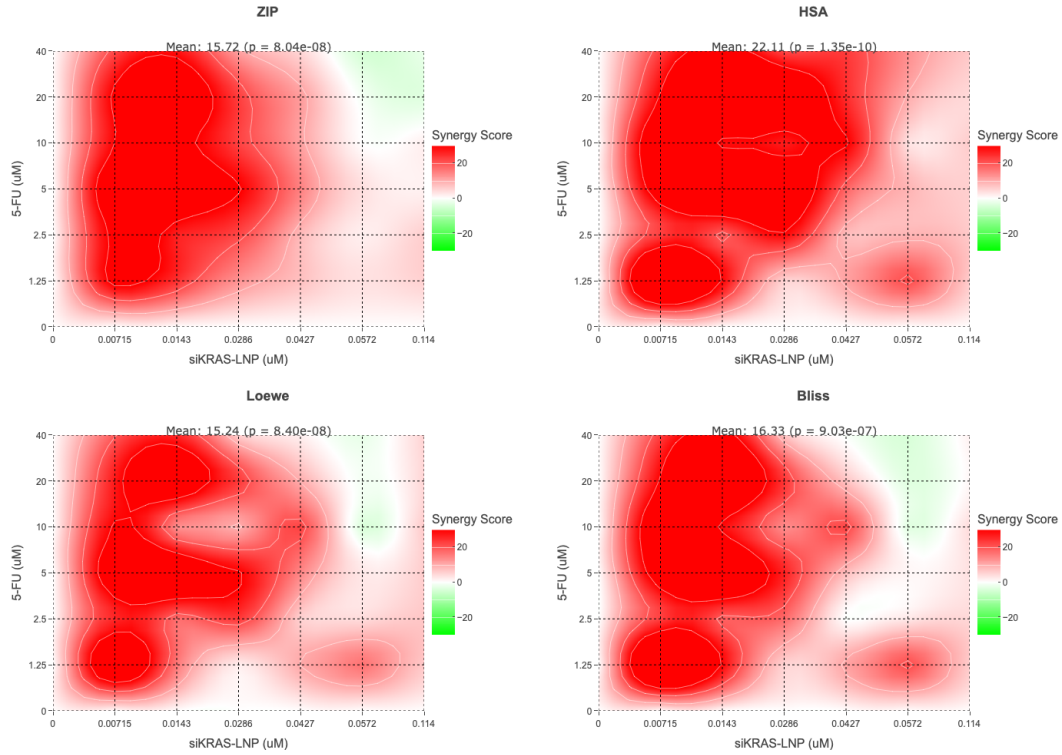

## LS180

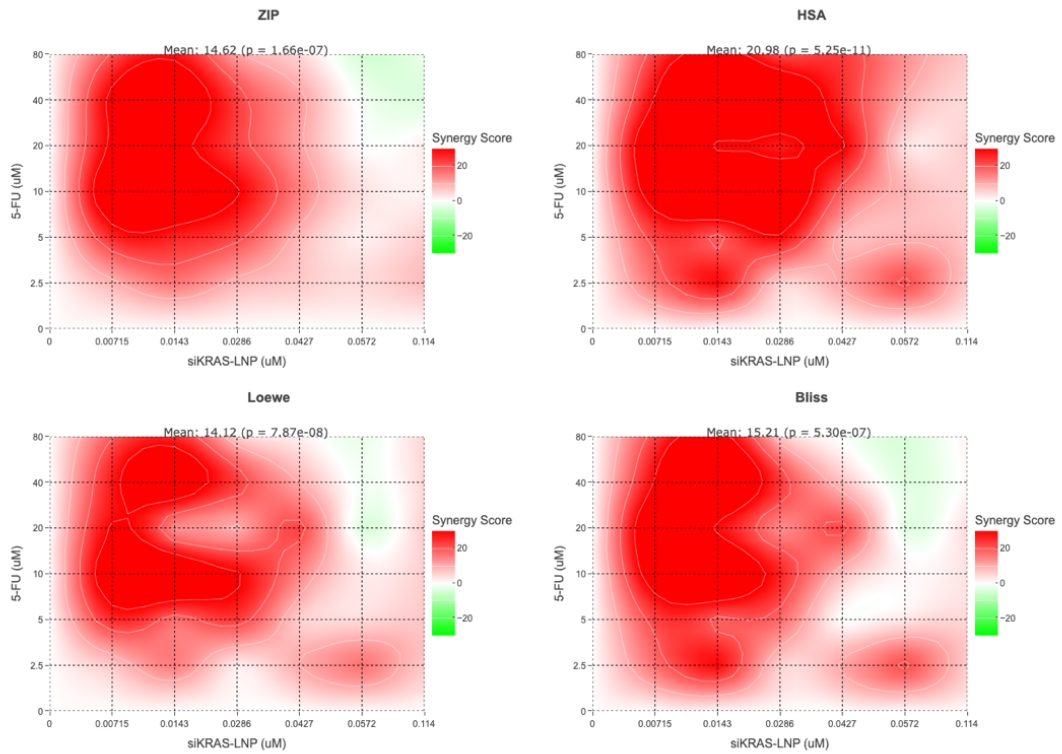

**Figure S10.** Synergy scores using the ZIP, HSA, Loewe, and Bliss reference models following combination treatment.
